# Regulatory remodeling in the allo-tetraploid frog *Xenopus laevis*

**DOI:** 10.1101/120212

**Authors:** Dei M. Elurbe, Sarita S. Paranjpe, Georgios Georgiou, Ila van Kruijsbergen, Ozren Bogdanovic, Romain Gibeaux, Rebecca Heald, Ryan Lister, Martijn A. Huynen, Simon J. van Heeringen, Gert Jan C. Veenstra

**Author notes:** Equal contributions (Dei M. Elurbe, Sarita S. Paranjpe, Georgios Georgiou).

## Abstract

**Background:** Genome duplication has played a pivotal role in the evolution of many eukaryotic lineages, including the vertebrates. The most recent vertebrate genome duplication is that in *Xenopus laevis*, resulting from the hybridization of two closely related species about 17 million years ago [1]. However, little is known about the consequences of this duplication at the level of the genome, the epigenome and gene expression.

**Results:** Of the parental subgenomes, S chromosomes have degraded faster than L chromosomes ever since the genome duplication and until the present day. Deletions appear to have the largest effect on pseudogene formation and loss of regulatory regions. Deleted regions are enriched for long DNA repeats and the flanking regions have high alignment scores, suggesting that non-allelic homologous recombination (NAHR) has played a significant role in the loss of DNA. To assess innovations in the *X. laevis* subgenomes we examined p300 (Ep300)-bound enhancer peaks that are unique to one subgenome and absent from *X. tropicalis*. A large majority of new enhancers are comprised of transposable elements. Finally, to dissect early and late events following interspecific hybridization, we examined the epigenome and the enhancer landscape in *X. tropicalis × X. laevis* hybrid embryos. Strikingly, young *X. tropicalis* DNA transposons are derepressed and recruit p300 in hybrid embryos.

**Conclusions:** The results show that erosion of *X. laevis* genes and functional regulatory elements is associated with repeats and NAHR, and furthermore that young repeats have also contributed to the p300-bound regulatory landscape following hybridization and whole genome duplication.

## Background

Genome duplication is a major force in genome evolution that not only doubles the genetic material but also facilitates morphological innovations. In plants whole genome duplications (WGD) appear to occur more often than in animals [2] and some phenotypic innovations, like the origin of flowers, have been attributed to this phenomenon [3]. In animals, two rounds of WGD at the root of the vertebrate tree (∼ 500 million years ago, Mya) gave rise to the four HOX clusters and have led to the expansion of the neural synapse proteome [4]. It is likely that this facilitated an increase in the morphological complexity [5] and allowed an increase in the complexity in the vertebrate behavioral repertoire [6]. More recent genome duplications have been documented in fish, at the root of the teleost fish 320 Mya and in the common ancestor of salmonids 80 Mya [7]. Amphibians in general appear to have undergone many polyploidisations, with natural polyploids in 15 Anuran and in 4 Urodelan families. In *Xenopus* (African clawed frogs) duplications have occurred on multiple occasions, giving rise to tetraploid, octoploid, and dodecaploid species [8]. The most recent documented duplication in vertebrates occurred in the amphibian *Xenopus laevis* 17 Mya [1]. The allo-tetraploid genome of *X. laevis* consists of two subgenomes, referred to as L (long chromosomes) and S (short chromosomes), that originated from distinct diploid progenitors [1]. Most of the additional genes that result from WGD events tend to be lost in evolution. In the case of allopolyploidy this loss is biased to one of the parental subgenomes [9], a phenomenon referred to as biased fractionation. One explanation for biased fractionation is the variation in the level of gene expression between the homeologous chromosomes [10], with the lowest expressed gene having the highest probability of being lost because it would contribute less to fitness.

The effects of polyploidization on the epigenome have mainly been studied in plants, where correlations between the gene expression and epigenetic modifications have been observed between homeologous genes [11], but are not well characterized in animals. The epigenetic modifications found in chromatin (DNA methylation and post-translational modifications of histones) are involved in gene regulation during development and differentiation [12, 13]. A high density of methylated CpG dinucleotides is repressive towards transcription; conversely, the DNA of a large fraction of promoters is unmethylated. In addition, histone H3 in promoter-associated nucleosomes is tri-methylated on lysine 4 (H3K4me3) when the promoter is active. Active enhancers on the other hand are decorated with mono-methylated H3K4 (H3K4me1) and they also recruit the p300 (Ep300) co-activator which can acetylate histones. When genes are expressed, they not only recruit RNA polymerase II (RNAPII), responsible for the production of the mRNA, but the gene body will be decorated with H3K36me3, which is left in the wake of elongating RNAPII. Therefore, deep sequencing approaches to determine these biochemical properties in a given tissue or developmental stage can be used to interrogate the activity of genomic elements. This is highly relevant in the context of genomic evolution, as changes in gene expression caused by mutations in *cis*-regulatory elements are a major source of morphological change during evolution [14].

Here we ask how genome evolution and the epigenetic control of gene expression are related to interspecific hybridization and whole genome duplication. We compare functional regulatory elements in the L and S subgenomes of *X. laevis* embryos by ChIP-sequencing of histone modifications, RNA-sequencing and whole genome bisulfite sequencing and use *Xenopus tropicalis*, a closely related diploid species as a reference. We quantify the loss and the gain of genetic material and analyze how it has affected genes and gene-regulatory regions. Although genome evolution after the hybridization appears dominated by sequence loss, we also find evidence for the gain of functional elements. We specifically identify new subgenome-specific regulatory elements that recruit p300 and show that these are enriched for transposable elements. Finally, to assess the early gene-regulatory effects of hybridization we analyze experimental interspecific *X. tropicalis × X. laevis* hybrids and we observe hybrid-specific p300 recruitment to DNA transposons, further highlighting the role of such elements in the evolution of gene regulation.

## Results

### The ***X. laevis*** L and S subgenomes show a bias in chromatin state and gene expression

To study the evolution of gene regulation in the context of whole-genome duplication we generated transcriptomic and epigenomic profiles in *X. laevis* early gastrula embryos (Nieuwkoop-Faber stage 10.5; Additional file 1). We performed RNA-sequencing and obtained epigenomic profiles using chromatin immunoprecipitation followed by deep sequencing (ChIP-seq). We generated ChIP-seq profiles for H3K4me3, associated with promoters of active genes, H3K36me3, associated with actively transcribed genes, the Polr2a subunit of RNA Polymerase II (RNAPII) and the transcription coactivator p300. In addition, we performed whole genome bisulfite sequencing to obtain DNA methylation profiles [15]. The sequencing results and details are summarized in additional file 1.

We created whole genome alignments (see Methods) to establish a framework for analysis of the epigenetic modifications in the two *X. laevis* subgenomes and in the *X. tropicalis* genome. Respectively 61% and 59% of the *X. laevis* L and S non-repetitive sequence can be aligned with the orthologous *X. tropicalis* sequence. This allows for comparisons of the activity of genes and regulatory elements between homeologous regions. Figure 1 shows a region on *X. tropicalis* chromosome 8 containing four genes, together with the corresponding aligning sequences on chr8L and chr8S in *X. laevis*. The epigenomic profiles (H3K4me3, p300, RNAPII and H3K36me3) of both *X. laevis* and *X. tropicalis* [16] are shown and the sequence conservation obtained from the whole gene alignment is illustrated by grey lines in the center of the plot. Regions that are conserved at both the sequence level and at the functional level (as measured by ChIP-seq) are highlighted. The *anp32e* gene is an example of a conserved gene that is expressed from all three genomes, as evidenced by H3K4me3 at the promoter and H3K36me3 and elongating RNAPII in the gene body. In contrast, expression of the *plekho1* gene has been lost from S. The gene is still present, but it is not active. There is no evidence of expression and both the H3K4me3 and the p300 signal are lost. Finally, the *vps45* gene is an example of a gene that is completely lost from L.

**Figure 1.**
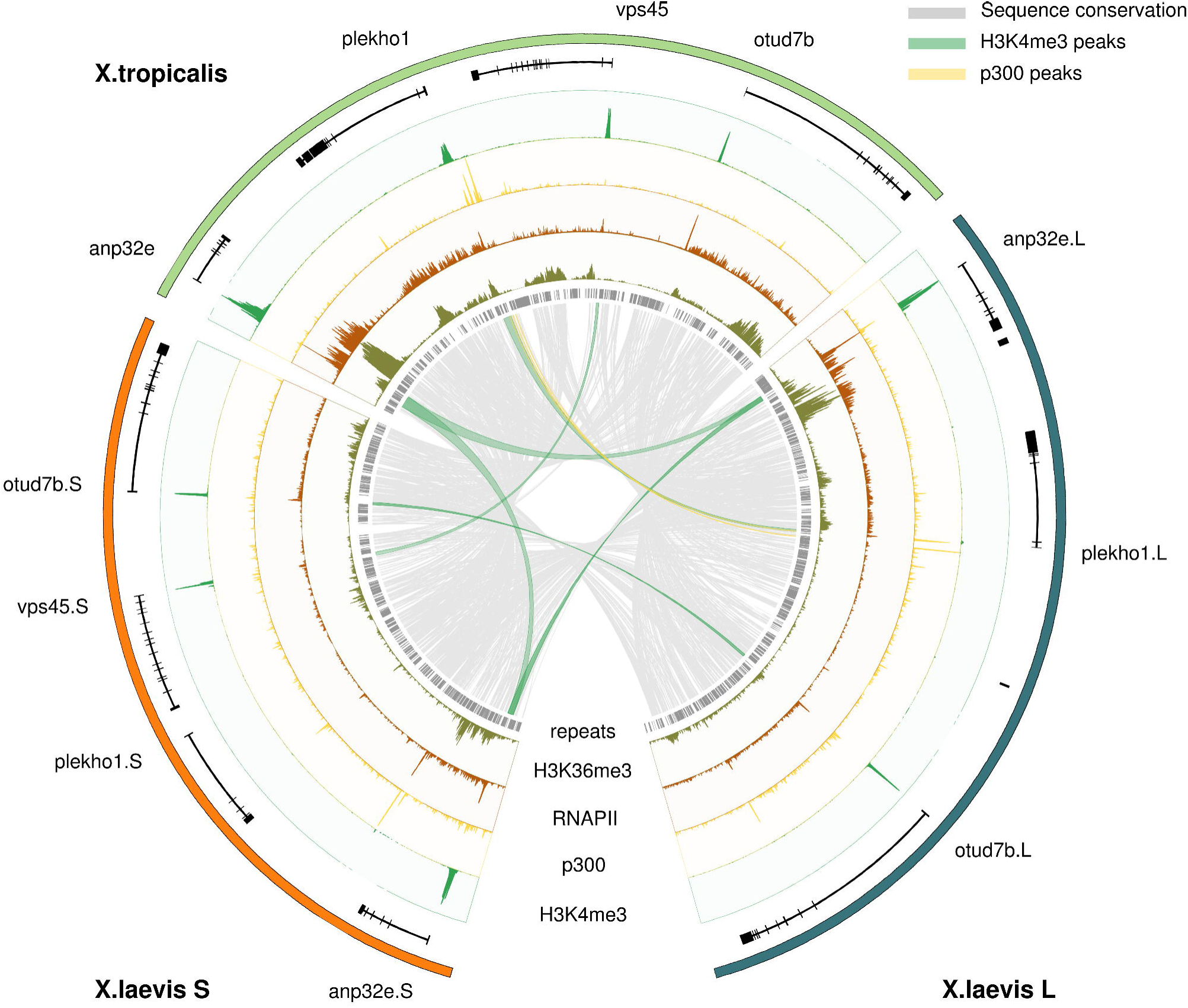
Alignment of a region on chromosome 8 in *X. tropicalis* and the *X. laevis* L and S subgenomes annotated with experimental ChIP-seq data (gastrula-stage embryos; NF stage 10.5). Shown are the gene annotation (black), repeats (grey), ChIP-seq profiles for H3K4me3 (green), p300 (yellow), RNA Polymerase II (RNAPII; brown) and H3K36me3 (dark green). The sequence conservation is indicated by grey lines. Conserved H3K4me3 and p300 peaks are denoted by green and yellow lines respectively. The anp32e gene is expressed in *X. tropicalis* and both the L and S subgenome of *X. laevis*. The *plekho1* gene, on the other hand, has lost promoter and enhancer activity on the *X. laevis* S locus, and shows no experimental evidence of being expressed.

Next, we quantified gene expression patterns in the *X. laevis* subgenomes. Of the 11,818 genes expressed at stage 10.5, 6,971 are located on L chromosomes and 4,847 on S. As reported previously [1], a minor but significant expression bias is detected among pairs of homeologs where both genes have detectable expression (Spearman R = 0.80; median expression difference of L compared to S = 5.7%; Fig. 2a). However, for many homeologs the expression bias is quite high, such that for one copy hardly any expression can be detected. Such non-expressed homeologs are located on both L and S, but occur more frequently on S (L: 41%, S: 59%).

**Figure 2.**
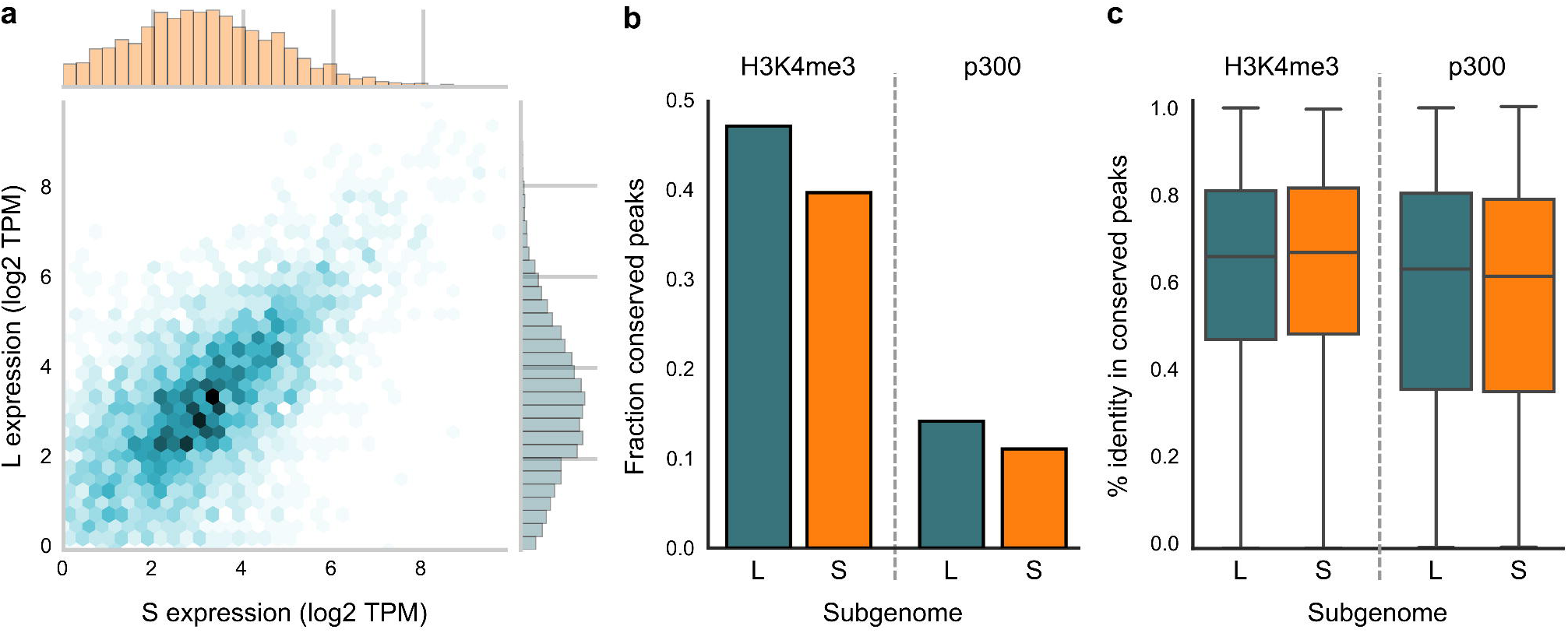
(a) Scatterplot of the expression level (log2 TPM) of L and S homeologs that are both expressed. The expression level of homeolog genes is generally similar (Spearman R 0.80). (b) Fraction of epigenetic signals (“peaks”) conserved in *X. laevis* compared to *X. tropicalis*. Promoters appear more conserved than enhancers; S has lost more epigenetic elements than L. (c) Active functional elements are equally conserved between L and S as compared to *X. tropicalis*. The background level of sequence conservation in fourfold degenerate sites from coding sequences with respect to *X. tropicalis* is 78.4% in L and 77.7% in S

We examined whether the expression differences between the L and S homeologs could be explained by differential transcription regulation. We used the epigenomic profiles to assay the promoter state (H3K4me3, DNA methylation), enhancer activity (p300) and active expression (RNAPII, H3K36me3). The L subgenome has 38% more annotated genes than the S subgenome [1]. We observe the same trend for the regulatory elements. The number of H3K4me3 peaks, DNA-methylation free regions (see Methods) and p300 peaks is higher on L (28%, 23% and 35%, respectively; Additional file 2). The overall effect is that there is no significant difference between the numbers of regulatory elements per gene for the two subgenomes.

To analyze the conservation of regulatory elements, we compared the H3K4me3 and p300 data to similar ChIP-seq profiles from *X. tropicalis* obtained at the equivalent developmental stage [16]. In general promoters are much more conserved than enhancers (Fig. 2b). From all H3K4me3 peaks in *X. tropicalis*, ∼40% are conserved in *X. laevis*, while for the p300 peaks the conservation is only ∼13%. This is congruent with the finding in mammals that enhancers evolve much more rapidly than promoters [17]. Whereas the number of conserved regulatory elements is lower in S than in L, the elements that can be aligned differ relatively little at the sequence level and show over ∼60% sequence identity (Fig. 2c).

These analyses show that the L and S subgenomes have evolved differently not only with respect to gene content [1], but also in terms of gene expression and chromatin state. Many more genes from S are lower expressed than their homeologs in L than vice versa. The number of functional regulatory elements, as identified by H3K4me3 and p300 ChIP-sequencing, is in line with a more profound loss of homeologous genes from the S subgenome. Next we set out to determine the origin of this differential loss.

### Large deletions are prominent in the S subgenome

The chromosomes of the *X. laevis* S subgenome are substantially shorter than the L chromosomes. The average size difference is 17.3% based on the assembled sequence [1] and 13.2% based on the karyotype [18]. To investigate the cause of these differences, we analyzed the pattern of deletions on both subgenomes. We called deleted regions based on the absence of conservation between the *X. laevis* subgenomes if they were at least partly conserved between one *X. laevis* subgenome and *X. tropicalis*. In addition, to be able to measure the size of the deletions, we required that the putative deleted regions were flanked on both sides by conserved sequences on both *X. laevis* subgenomes (Additional file 3: Figure S1). This resulted in a set of 19,109 deletions, of which 13,066 (68%) were deleted from S (LΔS) and 6,043 (32%) were deleted from L (SΔL). There is a clear deletion bias towards S, which increases with the size of the deletion (Fig. 3a). These deletions affect genes and their regulatory sequences, as for example in the *glrx2* locus where the promoter and most of the exons have been lost from the S subgenome (Fig. 3b). Overall, the bias in deletions towards S is larger when the regions are predicted to be functional. The fraction of sequence lost from intergenic regions in S is 2.3 times the fraction lost from L, in introns this deletion bias is 2.8-fold while in exons it is 4.6-fold (Fig. 3c, left). Interestingly, when we put deletions in an actively transcribed gene context, (IntronicTx, ExonicTx, see Methods: Active transcription), the deletion bias is again larger in exonic regions than in intronic regions but lower in general (Fig. 3c, center). Actively transcribed genes feature fewer deletions compared to all genes, which could be explained by the finding that highly expressed genes tend to be preferentially retained throughout evolution [19]. We also observe a high deletion bias for functional elements identified by ChIP: H3K4me3 (4.6 times) and p300 peaks (3.5 times) (Fig. 3c, right). These results substantiate that there is a deletion bias in S relative to L that is higher when the region appears to be functional. The differences in bias are mostly due to differences in the deletions from L (SΔL), suggesting that functional sequences on L have experienced more negative selection.

**Figure 3:**
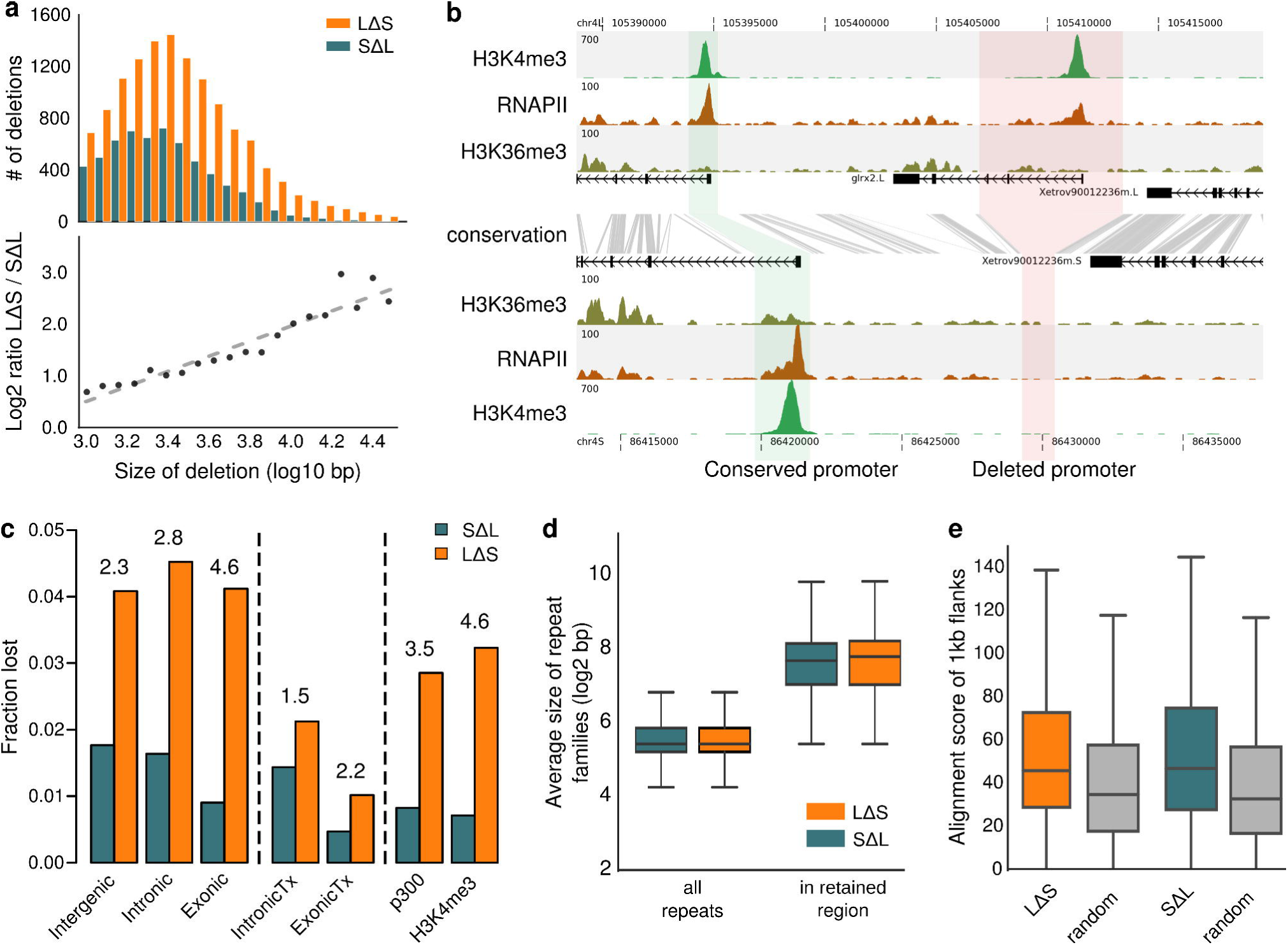
The S subgenome has more and larger deletions than L. (a) Size frequency distribution of deletions (top panel) and size ratio of L Delta S deletions relative to S Delta L deletions as a function of deletion size (bottom panel) (b) An example of a gene (*grlx2*) that has lost the promoter on the S genome due to a deletion. Shown are the gene annotation (black), ChIP-seq profiles for H3K4me3 (green), RNAPII (brown) and H3K36me3 (dark green). The sequence conservation is indicated by grey lines. (c) Fraction of genomic regions lost by deletions. Numbers on top of the bars represent the ratio of the fraction lost in S relative to the one lost in L. Intergenic: 1kb distance from a gene. Intronic: introns. Exonic: UTRs + CDS. IntronicTx: introns from genes actively transcribed. ExonicTx: Exons from genes actively transcribed. p300: genomic fragments having a p300 peak. H3K4me3: genomic fragments having a H3K4me3 peak. (d) Retained regions associated with deletions are enriched for relatively long repeats (e) 1kb flanks of the retained regions are more similar to each other than random genomic regions of the same size.

One of the possible sources of the loss of genomic DNA in the L and S subgenomes is non-allelic homologous recombination (NAHR), which is known to occur between long repetitive elements on the same chromosome [20]. To test whether this phenomenon could be responsible for the genomic losses detected, we examined the length distribution of repetitive elements in retained regions, i.e. the homeologous regions of the sequences that were lost in one of the subgenomes (Fig. 3d). Indeed, we observe that repetitive elements are on average 3.7 times longer (P< 1e-52, Mann-Whitney U) compared to random genomic sequences (Fig. 3d). Furthermore, the flanks of the retained regions (L for LΔS and S for SΔL, respectively) tend to be more similar to each other than random genomic sequences (Fig. 3e). Nevertheless, the current density of repetitive elements is similar in the L and S subgenomes (Additional file 3: Figure S2), indicating that repeat density alone does not cause biased sequence loss on S chromosomes. These observations suggest that NAHR of ancient repeats has played a significant role in the deletions of regions from both subgenomes, most profoundly from the S chromosomes. To estimate when in the evolution these deletions and other types of mutations occurred we dated the origin of the pseudogenes that they caused.

### High levels of pseudogenization started after hybridization and continue to the present

To date the pseudogenes, we aligned them with the protein coding regions in L, S and the outgroup *X. tropicalis* (Methods: Search and alignment of orthologs and evolution rates). The non-pseudogene coding regions in S are generally less conserved than in L, especially regarding synonymous substitutions (Ks, Fig. 4a) with the ratio between nonsynonymous and synonymous substitutions (Ka/Ks) only slightly higher in S than in L (Fig. 4b). The difference in Ks between the L and S subgenomes shows that S has been subject to moderately higher mutation rates than L. In order to examine whether the relatively high level of mutations in the S genome persists to this day, we examined the level of SNPs separating the published inbred genome [1] and the progeny of two outbred individuals (Methods: SNP calling). We observe that the level of SNPs in the S genome is consistently 3% higher than in the L genome irrespective of whether the SNPs are in intergenic regions, intronic regions or 4-fold degenerate positions of coding DNA (i.e., all the genomic regions assumed to be under relaxed constraint) (Additional file 4). The higher level of SNPs in coding DNA than in non-coding DNA correlates with an overrepresentation of CpGs in coding DNA (Additional file 3: Figure S3) and has been observed before in human genomes [21].

**Figure 4:**
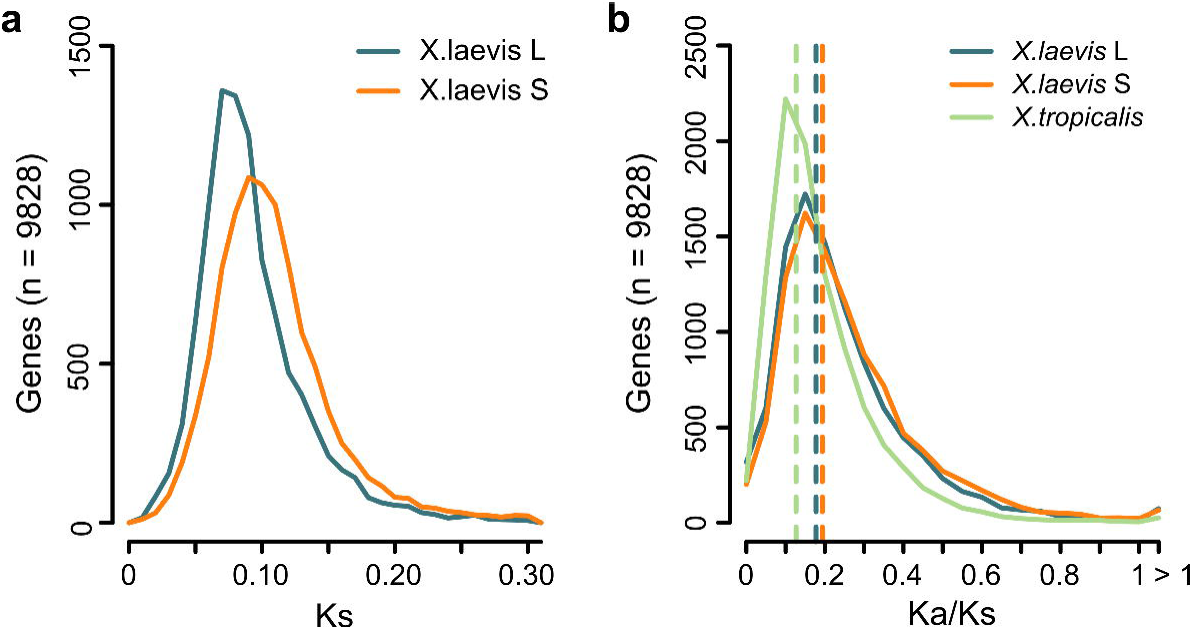
The S subgenome has a higher mutation rate than L. (a) Ks distribution per subgenome in *X. laevis*. (b) Ka/Ks in *X. laevis* and *X. tropicalis*.

The higher SNP density in S relative to L (Additional file 4) suggest that the relatively high rate of genome degradation in S continues to this day. To examine the continuity of this genome degradation we dated unitary pseudogenes [22] caused by point mutations and / or deletion-related events (Fig. 5a). We distinguish four, non-exclusive types of pseudogenes: genes that contain a premature stop codon, genes of which the coding sequence is at least 50% shorter than their homeolog and their ortholog in *X. tropicalis*, genes that have lost at least the 75% of their promoter relative to their homeologs that do have a promoter decorated with H3K4me3 in embryos, and genes that contain a frameshift. We furthermore required for each class that the pseudogene candidate is expressed at least 10-fold lower than its homeolog. In all cases, we do observe that the rate of pseudogenization has increased dramatically around 18 Mya, i.e. close to the inferred date of the hybridization, and that that rate is ∼2.3 fold higher in S than in L (Fig. 5a). Furthermore, this rate continues to be high until this day for every class considered (Fig. 5b). We obtained very similar results when we included one-to-one orthologs from additional species in the dating of the pseudogenes and bootstrapped the results per gene to obtain confidence intervals (Methods, Bootstrapping pseudogene dates) (Additional file 3: Figure S4). When we separate the pseudogenes into non-overlapping classes we observe that deletions are a prevalent cause of pseudogenization (39% and 44% on resp. L and S), and, as expected, the older pseudogenes are affected by more than one type of damage (Additional file 3: Figure S5). Pseudogenization after genome duplication has been observed to affect certain classes of protein functions more than others, with metabolic functions often being the first ones to be lost relative to regulatory proteins [7]. Indeed, when we date the loss of genes in the function categories associated with the loss, we find an overrepresentation of various metabolic processes, with the pseudogenes belonging to those categories dating often shortly after the WGD event (Additional file 3: Figure S6).

**Figure 5:**
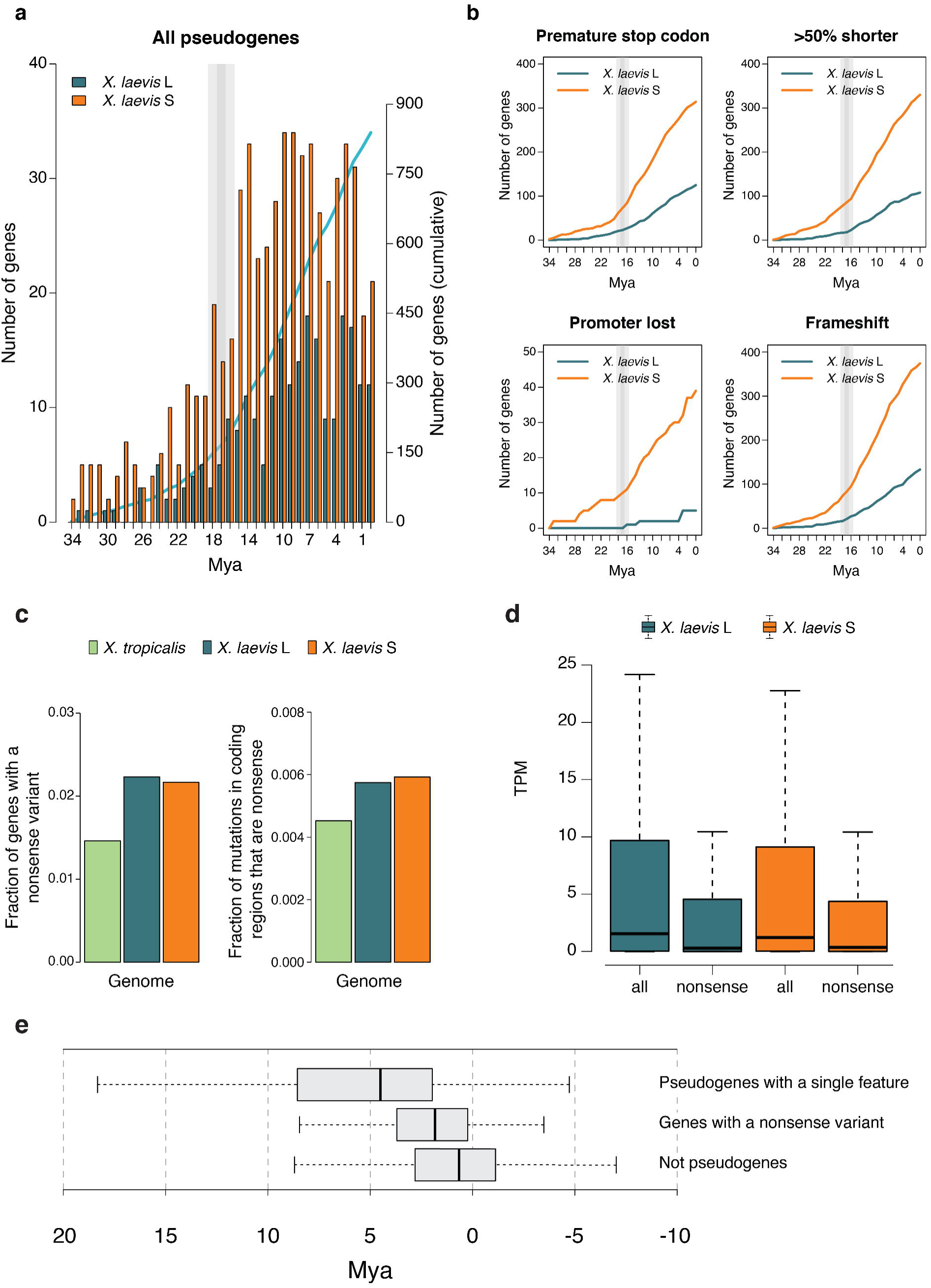
Pseudogenization rate has increased after hybridization. (a) Number of likely pseudogenes (i.e., genes having one or more pseudogene feature and no expression while their homeolog is expressed) binned by predicted date of pseudogenization event. (b) Pseudogenes with different (non-exclusive) pseudogene features and their sum over the years. (c, left) Fraction of genes that have a nonsense variant in the population. (c, right) Fraction of mutations in coding regions that introduce a premature stop codon. (d) Expression of genes with and without a nonsense variant present in the population. (e) Distribution of predicted pseudogenization time (including one-to-one orthologs of human, mouse and chicken) for genes with a single pseudogene feature and a 10-fold lower expression than the homeolog (top), for genes with a nonsense variant present in the population of *X. laevis* (middle) and for genes that do not present any feature for pseudogenization and whose expression is less than 2-fold different between homeologs (bottom).

To find independent evidence that the rate of pseudogenization in *X. laevis* remains high until the present we examined genes that appeared to be polymorphic with respect to their pseudogene state: i.e. we searched for protein truncating variants (PTVs) (variants which potentially disrupt protein-coding genes) in the progeny of two of our outbred genomes (Methods: SNP calling) relative to the published inbred genome [1]. Among all possible PTVs, we limited the analysis to SNPs that introduce a premature stop codon (nonsense mutations), as they can be called relatively reliably [23]. As a reference, we compared the nonsense SNP density with the one we measured in *X. tropicalis* using the same type of data and settings to call the SNPs: i.e. the progeny of two outbred genomes. In the 23,667 annotated genes in L and 16,939 in S we detect 528 (2.23%) and 367 (2.17%) genes with at least one loss of function variant. In contrast, in the 26,550 genes of *X. tropicalis* we detect only 388 (1.46%) loss of function variants (Fig. 5c, left). When normalizing the nonsense variants by the total number of SNPs in coding regions per (sub) genome, the fraction of premature stop variants in S (5.9×10^-3^) is slightly higher than that in L (5.7×10^-3^) while both are substantially and significantly higher than in *X. tropicalis* (4.5×10^-3^; p < 0.001 for both comparisons; Chi-squared, Fig. 5c, right). To substantiate that the selected PTVs are indeed hallmarks of incipient pseudogenes, we compared their expression with the expression of the other genes in their respective (sub)genome and found that genes with a SNP introducing a premature stop codon have a significantly lower expression (Fig. 5d). Second, we found that genes with this type of variants present in the population show evidence of loss of selection when compared to the set of genes that are not pseudogenes (p = 10^-05^; Student’s t-test, Fig. 5e), and that this loss of selection is more recent than for pseudogenes with only a single feature for pseudogenization that is fixed in the population (p = 5.6×10^-07^; Student’s *t*-test, Fig. 5e).

Altogether, these results suggest that the differences that the subgenomes present nowadays are, at least partly, due to the combination of a higher mutation rate and a more relaxed selective pressure in S, leading to a differential gene loss. This gene loss continues to be at a higher rate than in a closely related diploid species.

### Transposons have contributed subgenome-specific enhancer elements

The results described above document the pervasive loss and ongoing decay of coding and regulatory sequences after interspecific hybridization genome duplication. We next asked to what extent regulatory innovations have contributed to genomic evolution of this species. At many loci, the profile of p300 recruitment is remarkably different between L and S loci, with differences in both p300 peak intensity and number of peak regions across homeologous loci, for example in the *slc2a2* locus (Fig. 6a). We identified 2,451 subgenome-specific p300 peaks lacking any conservation with either the other subgenome or *X. tropicalis* (colloquially referred to as ‘new’ enhancers). There are similar numbers of these non-conserved subgenome-specific p300-bound elements in the L subgenome (1,214) and the S subgenome (1,237).

**Figure 6.**
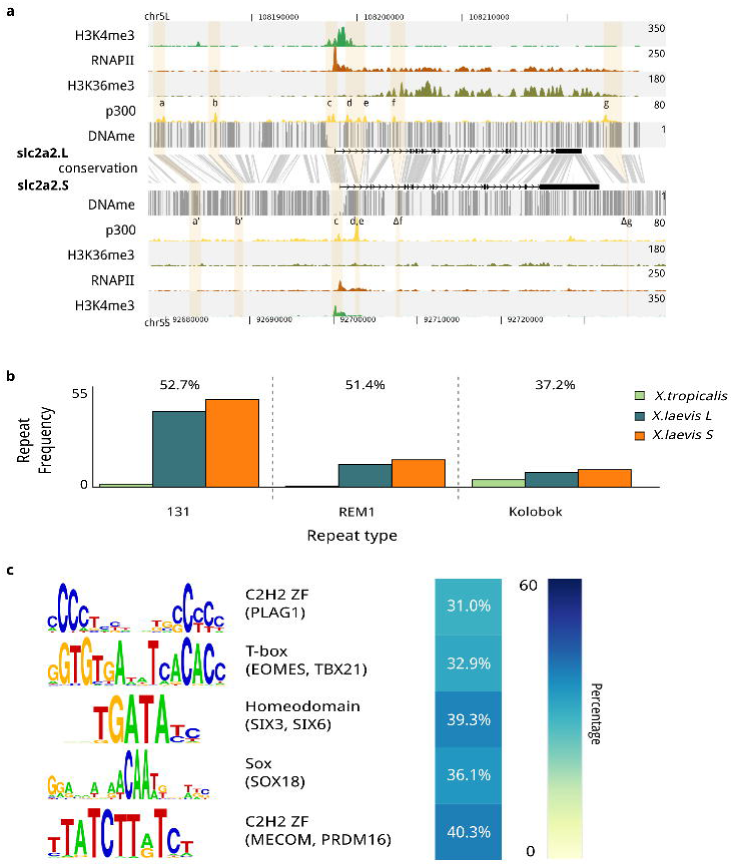
Sub-genome-specific recruitment of p300 is associated with transposable elements Subgenome-specific p300 peaks are enriched for transposable elements carrying transcription factor motifs active in early development. (a) Differential regulation of the slc2a2 homeologs at stage 10.5. Shown are the genomic profiles of H3K4me3 (green), RNA Polymerase II (RNAPII; purple), H3K36me3 (blue) and p300 (yellow) ChIP-seq tracks, as well as DNA methylation levels determined by whole-genome bisulfite sequencing (grey). The top panel shows slc2a2.L, which is highly expressed, as evidenced by RNAPII and H3K36me3, and has a number of active enhancers (a-g), while slc2a2.S, shown in the bottom panel, is expressed at a lower rate. The conservation between the L and S genomic sequence is shown in gray between the panels. Differential enhancers between L and S are highlighted in yellow, which illustrates lost enhancer function (a,b), conserved enhancer function (c-e) and deleted enhancers (f,g). (b) Subgenome-specific p300 peaks are associated with DNA transposon repeats (Threshold p-value ≤ 10^-4^, 2 times fold enrichment compared to all *X. laevis* peaks, and present at least in 15% of the peaks). The barplots show the frequency of occurrence of each of the three repeat types per megabase in the three (sub)genomes. Over the bars is represented the percentage of subgenome-specific peaks overlapping with the corresponding repeat. (c) Transcription factors found to be enriched in the subgenome-specific p300 peaks (Threshold p-value ≤ 10^-4^, 3 times fold enrichment compared to all *X. laevis* peaks, and present at least in 20% of the peaks).

Because new sequences can be acquired by transposition, we examined the overlap of subgenome-specific enhancers with annotated repeats and found that 87% (2,143 of 2,451) are associated with annotated repeats, three of which (designated REM1, Kolobok-T2 and family-131) were enriched in particular. The annotations individually overlap with 37-53% of the subgenome-specific p300 peaks, compared to 3-9% at other p300 peaks (Fig. 6b). Together these three annotations account for 1,338 (54%) of new enhancers, 862 of which have all three annotations overlapping at the same location. They form a 650 bp sequence with an almost perfect 195 bp terminal inverted repeat (TIR), the most terminal 65 bp of which shows 83-90% similarity with the TIRs of a Kolobok-family DNA transposon present in *X. tropicalis* (Additional file 3: Fig. S7). This specific Kolobok DNA transposon carries the REM1 interspersed repeat and is present almost exclusively in *X. laevis* (8,833 and 8,802 copies in resp. L and S, versus 4 copies in *X. tropicalis*), suggesting that it is a relatively young transposable element that proliferated after the split with *X. tropicalis*. It carries several transcription factor motifs, including the Eomes T-box motif and the Six3/Six6 homeobox motif (Fig. 6c).

We examined the correlation of the new Kolobok enhancers with gene expression and found that genes with a transcription start site within 5kb of these subgenome-specific Kolobok enhancers are more highly expressed than other genes in that subgenome (p = 10^-4^ for L and p = 8×10^-5^, Mann-Whitney U test) (Additional file 3: Fig. S8), suggesting that the new enhancers are inserted close to active genes and/or promote the expression of these genes.

### Regulatory remodeling by transposons in ***X. tropicalis × X. laevis*** hybrids

The gene expression (Fig. 2) and p300 recruitment (Fig. 6) differences between the L and S subgenomes may have been caused by regulatory incompatibilities affecting enhancer activity or DNA methylation, which could act immediately upon interspecific hybridization. Alternatively, these differences may represent the long-term effects of genomic co-evolution of the two subgenomes. To examine whether the differences between the two subgenomes were caused by the hybridization event itself, we determined the immediate effect of hybridization on DNA methylation and the patterns of H3K4me3 and p300 enrichment at regulatory regions. We generated embryos obtained by fertilization of *X. laevis* eggs (LE) with *X. tropicalis sperm* (TS). The resulting LETS hybrid embryos were compared to normal *laevis* (LELS) and *tropicalis* (TETS) embryos. The reverse hybrid (TELS) was not viable, as previously described [24].

To examine the early potential changes in DNA methylation, we performed whole genome bisulfite sequencing on the DNA of LETS, LELS and TETS embryos. The overall methylation in hybrid and normal embryos is almost identical at 92%. We identified a total of 709 differentially methylated regions (DMR) (FDR =0.05); 181 and 72 hypermethylated and 384 and 72 hypomethylated regions in respectively the *X. laevis* and *X. tropicalis* genomes. This reflects both gain and loss of DNA methylation in the sub-genomes of LETS hybrid embryos (Fig. 7f-g). There is no evidence in the underlying DNA sequence signatures for these regions being related to gene-regulatory regions (Additional file 3: Figure S9a-d). They are also not in close proximity of genes and may represent regions with inherently unstable DNA methylation. The global pattern of H3K4 trimethylation at promoters is also quite similar in hybrids and normal embryos; less than 10 peaks changed in hybrid embryos relative to normal embryos (Additional file 3: Figure S9e).

**Figure 7.**
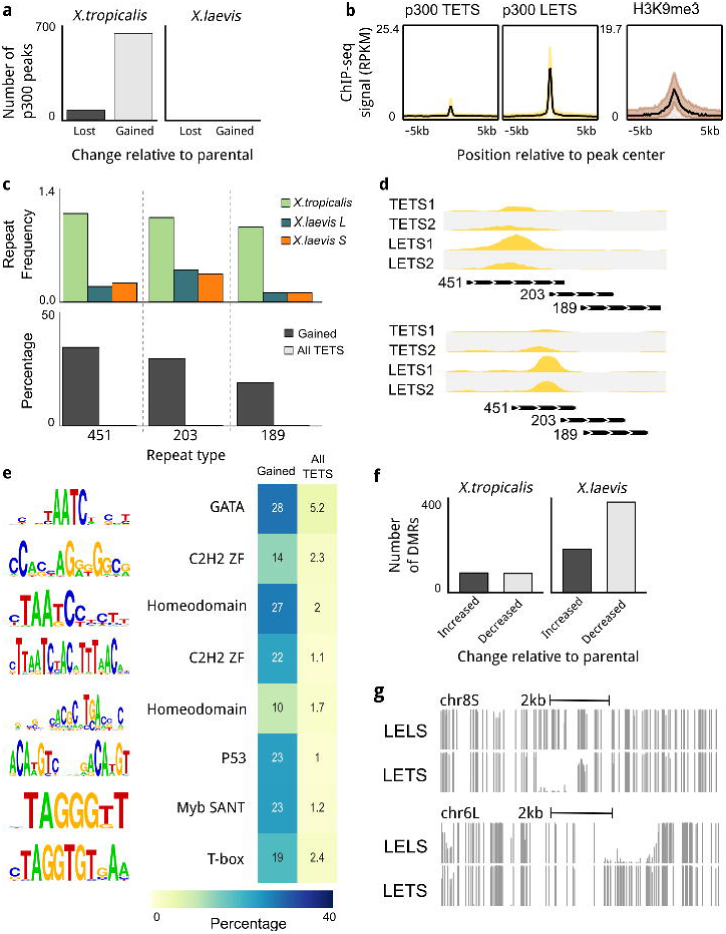
(a) Changes in p300 recruitment in LETS hybrids. In *X. tropicalis* genome there are new hybridization-induced peaks as well as peaks that disappeared after hybridization. In *X. laevis* genome there no changes. (b) Newly introduced peaks appear to be repressed by H3K9me3 in *X. tropicalis* embryos. (c, bottom) A significant number of hybrid-specific peaks are associated with DNA transposon repeats (Threshold p-value <= 10^-6^, > 20 times fold enrichment compared to all *X. tropicalis* peaks and present at least in 10% of the peaks). (c, top) The barplots show the frequency of occurrence of Motif:lcl|rnd-1_family-451_DNA, Motif:rnd-1_family-203 and Motif:lcl|rnd-1_family-189_DNA_PiggyBac repeats per megabase in the three (sub)genomes. Those repeats are *X. tropicalis* specific, as they occur more often compared to *X. laevis* genomes. (d) Profiles of *X. tropicalis* embryos p300 and LETS hybrid p300 in *X. tropicalis* hybridization-induced peaks loci. New peaks overlap with DNA transposons repeats. (e) Newly introduced peaks found to be enriched in TF DNA binding sites (Threshold p-value ≤ 10^-6^, 5 times fold enrichment compared to all *X. tropicalis* peaks, and present at least in 10% of the peaks). The transcription factors binding those motifs are Homeodomains, GATA, Zinc finger, P53, Myb SANT, Forkhead and T-box. (f) Differentially methylated regions (DMRs) in hybrid embryos (g) DNA methylation profiles showing the DNA methylation instability in LETS hybrids.

Recruitment sites of p300 however, are specifically gained and lost at several subsets of *X. tropicalis* genomic loci in hybrid embryos (Fig. 7a); 629 p300 recruitment sites were gained (a 2.6% increase relative to normal *X. tropicalis* embryos), whereas just 67 p300-bound regions were lost (adjusted p-value cutoff 1e-05). In the *X. laevis* part of the hybrid genome none were lost or gained (Fig. 7a), indicating that the changes in the hybrid are biased towards the paternal *tropicalis* genome. To assess the epigenetic state of the gained and lost p300-binding regions, we used our epigenome reference maps of histone modifications in *X. tropicalis* [16]. Among all the marks tested, only H3K9me3 was significantly enriched, specifically at sites of gained p300 recruitment (Fig. 7b), suggesting that these regions are heterochromatic in normal (TETS) embryos but can recruit the p300 co-activator in LETS hybrid embryos.

While examining the p300 hybrid-specific recruitment sites, we noticed that transposable elements were present at many locations (Fig. 7c, d); 82% of the hybrid-specific p300 peaks overlapped more than 50% with annotated repeats. We therefore examined the occurrence of specific repeats at gained p300 sites, and found that three repeat annotations (family-451, 203 and 189) were strongly enriched (p-value 1e-05; hypergeometric test), each accounting for 20-37% of all newly gained p300 peaks, whereas they only overlap with <1% of other p300 peaks (Fig. 7c, lower panel). The three repeat annotations strongly co-occur and form 1.3 kb repeat with a 200 bp imperfect TIR, which shows ∼80% similarity with those of known PiggyBac-N2A DNA transposons (Additional file 3: Figure S10). We recently found that DNA transposons that are heterochromatinized by H3K9me3 in *X. tropicalis* embryos are relatively young relative to other transposable elements [25]. Indeed, the piggyBac DNA transposons that gain p300 binding in hybrids are much less abundant in *X. laevis* than in *X. tropicalis*, suggesting that these relatively young transposons get derepressed in the *X. laevis* egg which has had little prior exposure to this transposon. These elements also carry transcription factor binding sites. Nine motifs are enriched (hypergeometric p-value 1e-05) and are present in 10-35% of gained p300 recruitment sites, compared to a 1-3% prevalence of these motifs in other p300 peaks (Fig. 7e). These DNA binding motifs represent binding sites of Homeodomain, GATA and T-box binding factors, which are abundantly expressed during early embryogenesis.

These results document DNA transposon-associated p300 recruitment and DNA methylation instability in experimental interspecific hybrids.

## Discussion

The genomes of the parental *Xenopus* species that gave rise to *X. laevis* through interspecific hybridization have remarkably been maintained as separate and recognizable subgenomes propagated on different sets of chromosomes [1]. These clearly distinguishable subgenomes allow detailed analyses of the patterns of (epi)genomic loss and regulatory remodeling. The loss of genes, regulatory elements and genomic sequence is caused predominantly by deletions and mutations in both subgenomes, which erode the S subgenome more strongly than the L subgenome. Such biased loss of genes has been observed in polyploid plant species and has been suggested to be a general result of allo-polyploidisation, in contrast to auto-polyploidies where the subgenomes are indistinguishable and degrade at a similar rate [9]. As to why one particular subgenome erodes more quickly than another, one hypothesis is that interspecific hybridization generates a crisis, referred to as ‘genomic shock’, for example by transposon reactivation on one of the subgenomes which can disrupt coding sequences [26]. Consistent with this possibility is the proliferation of S-specific Mariner DNA transposons in *X. laevis* at the time of hybridization [1]. Also consistent with transposon reactivation are our results from artificial *X. tropicalis × X. laevis* hybrids (LETS, *X. laevis* eggs, *X. tropicalis* sperm), in which a set of *X. tropicalis*-specific DNA transposons recruits the p300 co-activator in the hybrid, whereas normally they are repressed by H3K9me3.

Relatively young DNA transposons are heterochromatinized with H3K9me3 [25], but when introduced into eggs that have been little exposed to these transposons these mechanisms may fail. We have not been able to detect transposon expansion in the short time of *Xenopus* hybrid embryogenesis (data not shown), but together the observations suggest that transposon reactivation can contribute to genomic perturbations in hybrids. Similarly, in the Atlantic salmon, which has undergone several (320 MYA, 80 MYA) whole genome duplications, transposon expansion has been associated with the whole genome duplication event and with chromosome rearrangements [7].

In contrast to these short-term effects of hybridization, our analyses indicate that new pseudogenes continue to arise, both by mutations that cause premature stop codons, and by deletions that truncate the coding region or delete intergenic or promoter regulatory sequences. An elevated rate of pseudogene formation is observed on both the L and S subgenomes since the time of hybridization (∼17 Mya, cf. Fig. 5) up to the present day, suggesting genome erosion is a continuous process that has been and still is higher on S compared to L. Consistent with this result is a mildly elevated level of SNPs observed in S relative to L (Fig. 4; Additional file 4). The cause of the higher mutation rate of the S subgenome is unknown. The local mutation rate has been shown to correlate with replication timing [27], and it is possible that there are subtle but consistent differences in replication timing between the two subgenomes.

The higher level of genome degradation in S relative to L appears to be the result of a higher mutation and deletion rate in S, combined with less selection against the loss of (epi)genetic elements in S. The higher deletion and mutation rates are supported by higher numbers of deletions and mutations in regions that appear not to be under selection: intergenic regions, introns and redundant coding positions. Reduced selection against the loss of elements from S relative to L is supported by a higher deletion bias in p300 peaks, coding regions and promoters than in introns and intergenic regions, by a slightly higher Ka/Ks ratio in S and by a substantially larger amount of homeologs becoming pseudogenes in that subgenome. The deletions bear the hallmarks of NAHR [28]; the retained regions in the other subgenome are enriched for ancient repeats and the sequence similarity between the flanks of the region is higher than expected by chance. The S chromosomes have also experienced significantly more rearrangements including inversions [1]. Normally, in meiotic recombination double strand breaks are fixed using allelic sequences. In the absence of proper chromosome pairing, other non-allelic homologous sequences, for example repeats in the same chromosome, are used for double strand break repair, leading to deletions and inversions [28]. Interestingly, Prdm9, a fast-evolving mammalian DNA-binding protein involved in meiotic chromosome pairing and recombinational hotspot selection, has been implicated in hybrid sterility in mouse [29, 30]. There is no known one-to-one ortholog of Prdm9 in *Xenopus* and the L and S subgenome-encoded proteins involved in meiotic double strand break repair are also not fully known, but it is conceivable that their skewed expression or activity is involved in subgenome-biased NAHR.

The results reported here identify a major role for repetitive elements in subgenome bias, gene loss and regulatory remodeling. Not only sequence loss by NAHR is linked to repeats, subgenome-specific acquisition of enhancer elements is also overwhelmingly associated with transposable elements. Moreover, young transposons also gain p300 recruitment in *X. tropicalis × X. laevis* hybrids. DNA transposons can contribute sequence variation to the genome, which can affect gene expression by changing the local chromatin state at the site of insertion, resulting in metastable epi-alleles [26]. Once a host is invaded, transposable elements usually duplicate freely before they become repressed. When introduced in relatively unexposed eggs this repression may be lost. Interestingly, transposable elements can be co-opted as enhancers for the regulation of developmental genes [31, 32]. Transcription factors have been found to bind to transposable elements with open and active chromatin signatures in both human and mouse cells, but the binding patterns were largely different between the two species [33], suggesting that transposons contribute to regulatory change during evolution. In addition to the potentially large and sudden changes in regulatory potential caused by transposition, mutational changes are known to cause transcription factor binding sites to be lost and gained [17, 34], causing turnover and change in the regulatory landscape over longer time scales.

## Conclusions

It is not known exactly how the ancient two rounds of whole genome duplications at the root of the vertebrate tree have contributed to genome evolution. Its analysis is confounded by the pervasive loss of homeologs over hundreds of millions of years and the absence of tractable subgenomes. The *X. laevis* interspecific hybridization and genome duplication event is the most recent known vertebrate genome duplication. Excitingly, the clearly distinguishable chromosomes of different parental origins allow for reconstruction of the parental genomes. We have found evidence for a pervasive influence of repetitive elements, driving gene loss and genomic sequence loss through NAHR, in addition to remodeling of the regulatory landscape through transposon-mediated gain of coactivator recruitment. In combination with experimental interspecific hybrids, *Xenopus* can therefore be a powerful new model system to distinguish the short and long-term consequences of hybridization and to study the mechanisms of vertebrate genome evolution.

## Methods

### Animal procedures

Embryos were generated using IVF (*in vitro* fertilization) with outbred animals, including LELS embryos (laevis eggs-laevis sperm), TETS embryos (tropicalis eggs-tropicalis sperm) and LETS embryos (laevis eggs-tropicalis sperm). *X. laevis* female frogs were injected with 500U of hCG (human chorionic gonadotropin, BREVACTID 1500 I.E) 16 hours before IVF. A *X. laevis* male was sacrificed and isolated testis was macerated in 2 mL Marc’s Modified Ringer’s medium (MMR) to be used immediately for fertilization. Both male and female *X. tropicalis* frogs were primed with 100 and 15U of hCG 48 hours before IVF. Five hours prior to egg laying, females were boosted with 150U of hCG. Male testis was always isolated fresh. The testis was macerated in 2 mL FCS-L15 (10% fetal calf serum-90% L15 medium) cocktail and used immediately for IVF. LETS embryos were obtained similarly using species and sex-specific hormonal stimulation as described above. Once the macerated sperm suspension was mixed vigorously over the layered eggs, they were left undisturbed for three minutes and then the Petri dish was flooded with 25% MMR for the fertilized *X. laevis* eggs (LELS and LETS) and 10% MMR was added to the fertilized *X. tropicalis* eggs (TETS). Embryos were cultured at 25°C. The jelly coats were removed 4 hpf (hours post-fertilization) using 2% cysteine in 25% MMR (pH 8.0) for LELS and LETS and using 3% cysteine in 10% MMR (pH8.0) for TETS.

### ChIP-sequencing

Embryos (n = 35-90, two biological replicates for every ChIP experiment) were fixed in 1% formaldehyde for 30 minutes at Nieuwkoop-Faber stage 10.5. Embryos were washed once in 125 mM glycine / 25% Marc’s Modified Ringer’s medium (MMR) and twice in 25% MMR, homogenized on ice in sonication buffer (20 mM Tris·HCl, pH 8/10 mM KCl/1mM EDTA/10% glycerol/5 mM DTT/0.125% Nonidet P-40, and protease inhibitor cocktail (Roche)). Homogenized embryos were sonicated for 20 minutes using a Bioruptor sonicator (Diagenode). Sonicated extract was centrifuged at top speed in a cold table-top centrifuge and supernatants (ChIP extracts) were snap frozen in liquid nitrogen and stored at –20°C until use. Prior to assembling the ChIP reaction, the ChIP extract was diluted with IP buffer (50 mM Tris·HCl, pH 8/100 mM NaCl/2mM EDTA/1 mM DTT/1% Nonidet P-40, and protease inhibitor cocktail) and then incubated with 1–5 μg of antibody and 12.5 μl Prot A/G beads (Santa Cruz) for an overnight binding reaction on the rotating wheel in the cold room. The following antibodies were used: H3K4me3 (Abcam ab8580), H3K4me1 (Abcam ab8895), p300 (C-20, Santa Cruz sc-585), H3K36me3 (Abcam ab9050) and RNA polymerase II (Diagenode C15200004). The beads were sequentially washed, first with ChIP1 buffer (IP buffer plus 0.1% sodium deoxycholate), then ChIP2 buffer (ChIP1 buffer with 500 mM NaCl final concentration), then ChIP3 buffer (ChIP1 buffer with 250 mM LiCl), then again with ChIP1 buffer, and lastly with TE buffer (10 mM Tris, pH 8/1 mM EDTA). The material was eluted in 1% SDS in 0.1 M sodium bicarbonate. Cross-linking was reversed by adding 16 μl of 5 M NaCl and incubating at 65°C for 4–5 hours. DNA was extracted using the Qiagen QIAquick PCR purification kit. ∼ 10 ng input DNA was used for sample preparation for high-throughput sequencing on an Illumina HiSeq 2000 or NextSeq (according to manufacturer’s protocol).

### RNA-sequencing

For RNA-sequencing experiments total RNA was extracted from 20 Nieuwkoop-Faber stage 10.5 embryos (two biological replicates each for LELS and LETS respectively) using Trizol and Qiagen columns. 4-5 μg of total RNA was treated with DNase I on column and depleted of rRNA (ribosomal RNA) using Magnetic gold RiboZero RNA kit (Illumina) resulting in a yield of 45 - 52 ng of rRNA depleted total RNA. 2 ng of rRNA-depleted total RNA was reserved for Experion (Bio-Rad) quality assessment run for rRNA depletion and the remaining was used for first and second strand synthesis (strand-specific protocol). Total yield of dscDNA was between 14.5-15.8 ng and out of this 1.2 - 5 ng was used for sample preparation for high high-throughput sequencing (according to manufacturer’s protocol). qPCR quality controls before and after sample preparation corroborated well and relative depletion of 28S rRNA compared to control genes (eef1a1 and gs17) was taken as a quality assessment indicator for sequencing-grade dscDNA.

### ChIP-seq and RNA-seq data analysis

ChIP-seq reads were mapped to the *X. laevis* genome (Xenla9.1) using bwa mem (version 0.7.10-r789) with default settings [35]. Duplicate reads were marked using bamUtil v1.0.2. Where applicable (H3K4me3, p300) peaks were called using MACS (version 2.1.0.20140616) [36] relative to the Input track using the options ‐‐broad -g 2.3e9 -q 0.001. ‐‐buffer-size 1000. Peaks were combined for replicates using bedtools intersect (version v.2.20.1) [37]. Figures of genomic profiles were generated using fluff v1.62 [38]. In addition to the RNA-seq triplicate produced in this study, we used the eight stage 10.5 samples from NCBI GEO series GSE56586 (GSM1430926, GSM1430927, GSM1430928, GSM1430929, GSM1430930, GSM1430931, GSM1430932, GSM1430933). RNA-seq reads were mapped to the Xenla9.1 genome with the JGI 1.8 annotation using STAR version 2.4.2a [39]. Quantification of expression levels was performed using express eXpress version 1.5.1 [40]. The mean expression level (TPM; transcript per million) per transcript was obtained by combining all replicates.

### MethylC-seq for whole-genome bisulfite sequencing

Genomic DNA from *Xenopus* embryos (LELS and LETS, n = 20-50, NF stage 10.5) was extracted as described before [41] with minor modifications. Briefly, embryos were homogenized in 3 volumes STOP-buffer (15 mM EDTA, 10 mM Tris-HCl pH7.5, 1% SDS, 0.5 mg/mL proteinase K). The homogenate was incubated for 4 hours at 37°C. Two phenol:chloroform:isoamyl alcohol (PCI, 25:24:1) extractions were performed by adding 1 volume of PCI, rotating for 30 minutes at RT (room temperature) and spinning for 5 minutes at 13k rpm. DNA was precipitated in 1/5 volume NH4AC 4M plus 3 volumes EtOH with an overnight incubation at 4°C. Subsequently, the DNA was spun down for 20 minutes at 13k rpm in a cold centrifuge and the pellet was washed with 70% EtOH and dissolved in 100 μL of DNAse free water. To remove contaminating RNA, a 2 hours RNase A (0.01 volume of 10 mg/mL) treatment was performed at 37°C. Sample was further purified with two Mg/SDS precipitations. 0.05 volumes of 10% SDS plus 0.042 volumes of MgCl2 2M was added to the sample followed by incubation on ice for 15 minutes. Subsequently, the precipitants were spun down at 4°C for 5 minutes at 13k rpm. A third PCI extraction was also performed followed by only one chloroform:isoamyl alcohol (CI, 24:1) extraction. DNA was precipitated overnight at -20°C in 2.5 volumes EtOH plus 1/10 volume NaOAc 3M pH 5.2. Next, the precipitated DNA was spun down for 30 minutes at 13k rpm in a cold centrifuge and the pellet was washed with 70% EtOH. The purified DNA pellet was then dissolved in 50 μL H2O.

MethylC-seq library generation was performed as described previously [42, 43]. The genomic DNA was sonicated to an average size of 200 bp, purified and end-repaired followed by the ligation of methylated Illumina TruSeq sequencing adapters. Library amplification was performed with KAPA HiFi HotStart Uracil+ DNA polymerase (Kapa Biosystems, Woburn, MA), using 6 cycles of amplification. MethylC-seq libraries were sequenced in single-end mode on the Illumina HiSeq 1500 platform. The sequenced reads in FASTQ format were mapped to the in-silico bisulfite-converted *Xenopus laevis* reference genome (Xenla9.1) using the Bowtie alignment algorithm with the following parameters: -e 120 -l 20 -n 0 as previously reported [44, 45]. Differentially methylated regions were called using the methylpy pipeline, as described before [45], with FDR < 0.05 and the difference in fraction methylated larger than or equal to 0.4. To estimate the bisulfite non-conversion frequency, the frequency of all cytosine base-calls at reference cytosine positions in the lambda genome (unmethylated spike in control) was normalized by the total number of base-calls at reference cytosine positions in the lambda genome. See below for sequencing and conversion statistics.

DNA-methylation free (hypo-methylated) regions were detected using the hmr tool from MethPipe version 3.0.0 (http://smithlabresearch.org/software/methpipe/) [46]

### Active transcription

To consider a region as actively transcribed, we measured the H3K36me3 and RNAPII marks (as RPKM) of 200.000 random regions in *X. laevis*. We considered active transcription every region that showed at least the average of the measures plus two standard deviations, for both signals independently.

### Whole-genome alignment

Genome alignment of *X. tropicalis* and *X. laevis* was performed using progressiveCactus version 0.0 (https://github.com/glennhickey/progressiveCactus) [39, 40] with the default parameters. *X. tropicalis, X. laevis* L and S were treated as separate genomes and were aligned using (Xla.v91.L:0.2,Xla.v91.S:0.2):0.4,xt9:0.6) Newick format phylogenetic tree. In order to reduce computational time alignment was done per-chromosome, with homeologous chromosomes aligned to each other.

### Calling deletions

A set of high-confidence deleted regions was obtained using the progressiveCactus alignment. We extracted all regions from the *X. laevis* genome that reciprocally aligned either *X. tropicalis* and/or to the other subgenome. We then selected all regions that reciprocally aligned to *X. tropicalis* but not to the other *X. laevis* subgenome. We merged all regions within 10 bp and removed those that overlapped for more than 25% of their length with gaps. As a final filtering step, we required a sequence that reciprocally aligned to the other subgenome in both 500 bp flanks of the putative deletion. Finally, the size of the region between the two aligned flanks should be at most 4kb and at least 3 times shorter than the size of the region in the subgenome where the sequence was not deleted.

### SNP calling

SNPs were called using the GATK pipeline (version 3.4-46-gbc02625 [47]) on basis of the best practices workflow [48, 49]. As input we used a high-coverage ChIP-input track from a clutch of wild-type embryos compared the reference J-strain genome. The HaplotypeCaller tool was used to call SNPs. All putative SNPs were subsequently filtered with the VariantFiltration tool. The filterExpression was set to “QD < 2 || FS > 60.0 || MQ < 35.0 || MQRankSum < -12.5 || ReadPosRankSum < -8.0” for *X. tropicalis*. For *X. laevis* the same settings were used, except for MQ, which was set to “MQ < 40”. SNPs passing the filter were required to have at least ten-fold coverage with at least four observations of the alternative allele. The SNP coverage was calculated relative to the sequence regions where SNPs could be called given the minimum required coverage, as determined by the CallableLoci tool from the GATK pipeline.

### Search and alignment of orthologs and evolution rates

Orthologs of *X. tropicalis* were searched in the genome of*X. laevis* with the cdna2genome tool from Exonerate [50]. From 14500 sequences submitted, 14276 were successfully scanned. From those, 10935 found a match in both subgenomes, leaving 3343 sequences that did not return any sequence from either L or S subgenomes or both. Among the sequences with a match in both subgenomes, those having no synteny (939) were discarded because they were potential wrong matches in closely related gene families. Once we had our three sequences per gene (9996), we aligned them using MACSE [51], which allows frameshifts and premature stop codons, with the following parameters: gap creation -18, gap extension -8, frameshift creation -28, premature stop codon -50. 10 sequences were discarded in this step.

In order to obtain evolutionary rates of each of the three copies per gene triangle, we performed ancestral sequence reconstruction with FastML [52], which gave us the most likely sequence present at the speciation between *X. laevis* L and S ancestors. Once we obtained this crossroad sequence, we measured the amount of ratio of nonsynonymous mutations per nonsynonymous sites versus synonymous mutations per synonymous sites (i.e., Ka/Ks ratio) using the seqinR package [53].

### Pseudogene dating

Similar to Zhang et al. [22] we related the excess of nonsynonymous mutations to the evolving rate average of the gene to date the approximate time when the copy lost constraint on its sequence.

### Bootstrapping pseudogene dates

We took the pseudogene candidates and retrieved their annotated 1 to 1 orthologs in human, mouse and chicken through Ensembl. We then aligned them using MACSE [51] with default parameters, considering the pseudogene as a “less reliable” sequence. After this, we reconstructed the ancestral sequence with FastML [52] and then measured the Ka/Ks ratio using the seqinR package [53].

In order to confirm the reliability of these results, we bootstrapped the alignments 1000 times each and measured the Ka/Ks ratios of all of them. Briefly, we cut up the alignments in codons and we built an artificial alignment of the same length of the original protein by randomly adding (with replacement) aligned codons found in the original alignment.

### Quantification of genomic losses per genomic region

Using the deletions track generated through the deletions call step (see Methods: Calling deletions), we quantified the amount of DNA lost per genomic region by measuring the overlap between both coordinates. To do so, we used the R packages rtracklayer [54] and GenomicRanges [55].

### Gene Ontology term enrichment analysis

Term enrichment analysis was performed using PANTHER [56]. Briefly, we used *X. tropicalis* orthologs names of the pseudogenes discussed in section 6 and we compared it to the list of genes in *X. tropicalis* that successfully returned syntenic orthologs in *X. laevis* (see Methods: Search and alignment of orthologs and evolution rates).

## Declarations

### Acknowledgements

The authors thank Ulrike J. Jacobi and Kees-Jan François for valuable contributions in an early phase of the work and Emese Gazdag for genomic DNA.

### Availability of data and materials

The data have been deposited in NCBI’s Gene Expression Omnibus [34] and are accessible through GEO Series accession numbers GSE76059 (*X. laevis* ChIP-seq), GSE76059 (genomic DNA; *X. laevis* RNA-seq; *X. tropicalis x X. laevis* ChIP-seq), GSE90898 (X. *tropicalis × X. laevis* whole-genome bisulfite sequencing) and GSE67974 (*X. tropicalis* ChIP-seq)

### Authors’ contributions

ChIP-seq, RNA-seq data generation and experimental design was performed by SSP with help from IvK, RG and RH. GJCV, SJvH and MAH designed the study. DME and GG were involved in analysis design. Bisulphite sample generation and sequencing was done by SSP, IvK and OB, RL. OB and GG performed analysis of differentially methylated regions. Genome alignment and hybrid analysis was performed by GG. Analysis of deleted regions and SNPs was performed by SJvH and DME. DME also performed analysis of mutation rates and pseudogenes. DME, SSP, GG, MAH, SJvH and GJCV wrote the paper. DME, SSP, GG contributed equally to the study. All authors discussed the results and commented on the manuscript.

### Consent for publication

Not applicable

### Ethics approval and consent to participate

Not applicable

